# Development of a novel alpha7-nicotinic acetylcholine receptor-selective cell-penetrating peptide for intracellular cargo transport

**DOI:** 10.1101/2025.08.08.669423

**Authors:** Lahra Weber, Brittany C. V. O’Brien, Maegan M. Weltzin

**Affiliations:** Department of Chemistry and Biochemistry, University of Alaska Fairbanks, Fairbanks, AK, United States

**Keywords:** Cell penetrating peptide, drug delivery, biomimetic, alpha 7, nicotinic acetylcholine receptor, target selectivity, intracellular delivery, cargo delivery, rabies virus glycoprotein, α- bungarotoxin

## Abstract

Drug delivery into the brain remains a critical barrier for the treatment of neurological diseases. While the brain is shielded from many toxins and viruses by the blood-brain barrier (BBB), therapeutics are being created that exploit natural bypass mechanisms by forming complexes with cell-penetrating peptides (CPPs) derived from viruses. Neurological diseases often impact specific proteins in the brain; however, current CPPs lack the ability to selectively target precise cellular macromolecules. As a result, they are distributed broadly throughout the brain and cause off-target side effects. Neurotropic CPPs derived from the rabies virus glycoprotein (RVG) can access the brain by binding to plasma membrane targets, including, but not exclusively, nicotinic acetylcholine receptors (nAChRs). To overcome this barrier of minimal target selectivity, we designed several chimeric peptides composed of regions from the RVG and α-bungarotoxin, an α7 subtype-selective protein. Using human nAChRs expressed in *Xenopus laevis* oocytes, we screened the selectivity of our peptides using two-electrode voltage clamp electrophysiology. We identified a peptide with improved α7 nAChR subtype selectivity and apparent potency compared to the control RVG peptide. Using mammalian Neuro-2a cells, we demonstrated that our peptide depends on α7 nAChR plasma membrane expression to internalize and carry small-molecule payloads into the neuronal-like cells, without significant cytotoxic effects. Our novel α7 nAChR subtype-selective CPP may be useful in research applications requiring cargo delivery. Translationally, our α7 nAChR selective CPP holds potential to be a dual drug delivery system to transport cargo into the brain for the treatment of neurological diseases.

## Introduction

Delivery of therapeutics into the central nervous system (CNS) is often stymied by the blood-brain barrier (BBB). The BBB is a protective semi-permeable structure that restricts peripheral solutes, including drug molecules, from entering the brain, limiting the treatment of CNS disorders. Different approaches have been employed to overcome this barrier, including focused ultrasound disruption or intracerebral injections, to deliver therapeutics ^1–4^. While these methods may prove efficacious, they are also mild to moderately invasive, can require extended patient care time, or cause inflammation among other adverse effects ^3,4^. One approach to avoid these therapeutic pitfalls is the development of cell-penetrating peptides (CPPs) as drug delivery systems. These peptides are often non-toxic, and in some cases, facilitate the passage of payloads like nanoparticles, silencing RNA (siRNA), DNA, and other compounds across the BBB and into cells of the nervous system ^5–7^. However, CPPs generally interact broadly with cellular macromolecules and lipid bilayers, resulting in uncontrolled cargo delivery, undesired side effects, and ultimately failing to alleviate the disease burden. Engineering target-selective CPPs would eliminate many of these pitfalls while maintaining the beneficial therapeutic applications for CNS-targeted drug delivery.

One specific approach for the development of CPPs is the use of viral glycoprotein components, as these interact with host targets expressed on cell membranes to facilitate virion endocytosis ^6,8–11^. To exploit the neurotropic ability of the rabies virus, small CPPs have been created from its glycoprotein, and have been used to deliver functional cargo into the murine brain within 24hr ^12–18^. While this development is an important stepping stone for drug delivery into the brain, CPPs derived from the rabies virus glycoprotein so far lack target specificity ^19^. The rabies virus glycoprotein is known to interact with host proteins, including the neural cell adhesion molecule ^20–22^, p75 neurotrophin receptor ^23,24^, metabotropic glutamate receptor subtype 2 ^25^, integrin β1 ^26^, and nicotinic acetylcholine receptors (nAChRs) ^19,27,28^. We have recently demonstrated that a rabies virus glycoprotein-derived peptide (RVG) similar in sequence to that of other RVG CPPs ^12,14^ inhibits the function of the α7 nAChR by a competitive mechanism, in addition to heteromeric subtypes, albeit less efficaciously ^19^. These data demonstrated that RVG CPPs have functional actions independent of the attached cargo, and therefore, these need to be considered for therapeutic applications.

nAChRs are pentameric ligand-gated ion channels that initially activate in response to the neurotransmitter acetylcholine (ACh) to mediate cholinergic tone within the brain. Through combinations of the neuronal α2 - 7 and β2 - 4 subunits, a diverse population of homomeric and heteromeric subtypes are expressed in the CNS ^29^, with the α4β2 and α7 subtypes having the greatest abundance in the mammalian brain ^30,31^. The hippocampus and prefrontal cortex, areas of the brain responsible for learning, memory, and attention, are enriched with α7 nAChRs ^32,33^. Dysregulation or dysfunction of this subtype is associated with neurological diseases, including Alzheimer’s disease, Parkinson’s disease, schizophrenia, nicotine addiction, and major depressive disorder, making this subtype a therapeutic target ^34–39^. High affinity and nAChR subtype selectivity are obtainable with naturally occurring peptides found in the venom of animals ^40–45^. α-bungarotoxin (α-btx) is a venom component from the elapid Taiwanese banded krait snake (*Bungarus multicinctus)*. This ‘three-fingered protein’ has three loops, with loop 2 (lp2) interacting predominantly with the α7 nAChR orthosteric binding pocket. Devising chimeric peptides that utilize the existing high-affinity α-btx and RVG CPP may be a new avenue for increasing target selectivity.

We sought to harness the endogenous selectivity of the snake toxin α-btx with components of the rabies virus glycoprotein that have been used previously as a CPP, especially amino acid residues 174 - 203 (RVG), to generate an α7 nAChR-selective CPP. To accomplish this, we developed and screened a series of chimeric peptides on 10 nAChR subtypes and isoforms to evaluate selectivity using two-electrode voltage clamp (TEVC) electrophysiology. The cytotoxicity profile, cargo delivery, target selectivity, and CPP capabilities were assessed using the mammalian N2a cell line, which has a neuronal phenotype. Our results demonstrate that we have engineered a novel α7 nAChR selective CPP, a first of its kind, that has concentration-dependent antagonist activities, is not cytotoxic, and can transport cargo into neuronal-like cells. Clinically, our target selective CPP may be advantageous as it could be used as a dual-acting drug delivery system to transport therapeutic cargo into the brain for the treatment of neurological conditions involving the α7 nAChR subtype.

## Materials and Methods

### Reagents

Acetylcholine chloride (ACh), atropine sulfate, bovine serum albumin (BSA), dimethylsulfoxide (DMSO), Mecamylamine (Meca), and methyllycaconitine (MLA) were purchased from Sigma Aldrich (St. Louis, MO). All other buffer reagents were purchased from Sigma Aldrich unless otherwise specified. Chimeric peptides were designed by M. M. Weltzin (see Table 1 for sequences) and purchased from Elim BioPharmaceuticals (Hayward, CA) with HPLC purity > 90%. Lyophilized peptides were stored in the dark at -20°C until use. Peptides were first dissolved in DMSO before dilution with appropriate media or buffer. Drug solutions were made fresh daily and diluted as required.

**Table 1.** Sequence alignment of the RVG peptide and α-btx lp2 peptide. Bolded residues indicate sequence homology. Each peptide region (R) was tested and used to create the chimeric peptides.

| Virus/Toxin | Region 1 | Region 2 | Region 3 |
| --- | --- | --- | --- |
| RVG | **Y** T - I **W** | M P E N P R L G T S | **C D I F T N S R G K R A S K G** 203 |
| α-btx | **Y** R K M **W** | - - - - - - - - - - | **C D A F C S S R G K V V E L G** 43 |
| RRA | **Y** T - I **W** | M P E N P R L G T S | **C D A F C S S R G K V V E L G** 29 |
| ARA | **Y** R K M **W** | M P E N P R L G T S | **C D A F C S S R G K V V E L G** 30 |
| ARR | **Y** R K M **W** | M P E N P R L G T S | **C D I F T N S R G K R A S K G** 30 |

Eagle’s Minimum Essential Medium (EMEM), fetal bovine serum (FBS), Penicillin-Streptomycin, and pH 7.4 phosphate-buffered saline (PBS) were purchased from VWR International, LLC (Radnor, PA). Both Gibco Opti-Minimal Essential Medium (Opti-MEM) and Trypsin-EDTA (0.25%) were purchased from Gibco ThermoFisher Scientific (Waltham, MA). Lipofectamine 2000 Transfection Reagent and alamarBlue Cell Viability Reagent were purchased from Invitrogen (ThermoFisher Scientific).

### DNA constructs and cRNA synthesis

As described in our previous works ^19,46^, individual human nAChR subunits (α3 (NM_000743.5), α4 (NM_000744.5), α5 (NM_000745.3), β2 (NM_000748.2), β4 (NM_000750.5), and α7 (NM_000746.3) were generous gifts from Drs. Ronald J. Lukas and Paul Whiteaker (Barrow Neurological Institute, Phoenix, AZ). Also received was an α6/α3 chimeric subunit, which is comprised of the α6 extracellular domain and the α3 transmembrane and intracellular domain, and the linked subunit concatenated receptor β3α6β2α4β2. The α6/α3 chimeric subunit has increased expression compared to naturally occurring α6 subunits, while maintaining α6-like pharmacology ^47^. The β3α6β2α4β2 receptor was concatenated using 6 - 12 alanine, glycine, and serine repeats to link the subunits together and therefore enforcing the subunit positioning ^48^. Each nAChR subunit cDNA was received in the mammalian expression vector pCI (Promega, Madison, WI) except the α7 nAChR, which was in pSHE (a modified pGEMHE vector) and the concatenated receptor in pSGEM. The α7(345 - 348A) nAChR construct has residues 345 - 348 mutated to alanine, which abolishes interactions with Gαq and Gβγ without eliminating receptor expression or α-btx binding ^49^. α7(345 - 348A) DNA was synthesized from Twist Bioscience (San Francisco, CA) and inserted into the pSHE vector using XbaI and EcoRV restriction enzymes. cDNA amplification was accomplished via transformation with NEB® 5-α competent *E. coli* cells (New England Biolabs [NEB], Ipswich, MA) and extracted using the Qiagen QIAprep Spin Miniprep kits (Valencia, CA). Extracted DNA was verified via restriction enzyme digest. Constructs in the pCI vector were verified with enzymes XbaI and NotI, while constructs in pSHE/pSGEM were confirmed using XbaI and EcoRV. Digested DNA samples were visualized on a 1% agarose gel stained with ethidium bromide.

Before cRNA synthesis, cDNAs were linearized using restriction enzymes (SwaI for pCI and NheI for pSHE/pSGEM constructs). Linearized DNA was then treated with proteinase K (NEB) and purified using the Qiagen PCR clean-up kit. The T7 mMESSAGE mMACHINE™ High Yield Capped RNA Transcription Kit (Ambion, Austin, TX) was used to transcribe cRNA. Dual methods were employed to verify cRNA purity, including quantification by the Thermo Scientific NanoDrop 2000 spectrophotometer and electrophoresis gel imaging. cRNAs for each subunit or concatenated construct were sub-aliquoted and stored at -80°C.

The α7 pcDNA 3.1(+) and NACHO plasmid DNAs were prepared and verified as previously described in O’Brien et al. (2023) ^46^. The chaperone protein NACHO DNA plasmid in pREP9 was generously gifted by Dr. R. Loring (Northeastern University, Boston, MA). To facilitate robust expression of α7 nAChRs in the mammalian N2a cell line, the α7 subunit DNA was removed from the pSHE vector and subcloned into pcDNA 3.1(+). The final DNA construct was verified using XbaI, NotI, and PvuI restriction enzymes (New England BioLabs Inc., Ipswich, MA). To achieve transfection-grade, endotoxin-free constructs suitable for cell culture experiments, α7 nAChR and NACHO plasmid DNAs were extracted from 10-beta competent *E. coli* cells (High Efficiency; New England BioLabs Inc.) grown in Circle Grow broth media (MP Biomedicals, Santa Ana, CA) supplemented with 0.1 mg/mL ampicillin using the EndoFree Plasmid Maxi Kit (Qiagen, Valencia, CA). cDNA concentrations and purity (260/280 ratio) were measured using the NanoDrop 2000 spectrophotometer. cDNAs were kept frozen at -20°C until use.

### Oocyte preparation and cRNA injection

Expression of nAChRs was accomplished via cRNA injections into stage IV and V *Xenopus laevis* oocytes (Ecocyte Bioscience, Austin, TX). All efforts were made to minimize animal suffering and to reduce the number of animals used. Homomeric α7 and concatenated β3α6β2α4β2 nAChRs were expressed by microinjection of a single cRNA transcript (40 ng/oocyte). All other heteromeric nAChR subtypes were expressed via injection of mixed subunit cRNA (in ng) ratios (α4β2α5- 2.5 α4 : 2.5 β2: 25 α5; α6/α3β2β3- 12 α6/α3 : 12 β2: 6 β3; (α4β2)2α4-12.5 α4 : 0.125 β2; (α3β2)2α3 and (α3β4)2α3- 30 α: 1 β; (α3β2)2β2 and (α3β4)2β4- 1 α : 30 β. Each oocyte received a total of 81 nL of cRNA via impalement using a pulled glass micropipette (Drummond Scientific Company, Broomall, PA) with an outer diameter of ∼40 µm. Oocytes were incubated in buffer (82.5 mM NaCl, 2.5 mM KCl, 1 mM MgCl2, 1 mM CaCl2, 1 mM Na2HPO4, 5 mM HEPES, 600 μM theophylline, 2.5 mM sodium pyruvate, 50 μg/mL each penicillin, streptomycin, neomycin, gentamycin sulfate, pH to 7.5 using NaOH) at 13°C for 72 - 120hr prior to recording, with daily buffer changes.

### Electrophysiology

nAChR function was evaluated using two-electrode voltage clamp (TEVC) electrophysiology. No less than 72hr post-cRNA injection, *Xenopus laevis* oocytes were voltage clamped at -70 mV using an Axoclamp 900A amplifier with pClamp 10.6 software (Molecular Devices, LLC, Sunnyvale, CA) for data acquisition and analysis. DC offset was accomplished using a 40 Hz high-pass filter and a 10 kHz low-pass Bessel filter. Recording electrodes were pulled from thin-wall capillary glass (World Precision Instruments, Sarasota, FL) and filled with 3 M KCl. Electrode resistance ranged from 0.5 - 10 MΩ. Oocytes with leak currents below -100 nA were discarded.

Using a 16-channel, gravity-fed, perfusion system with automated valve control (AutoMate Scientific, Inc., Berkeley, CA), drug solutions were applied to voltage-clamped oocytes. Drug solutions were made in an oocyte ringer 2 (OR2) recording buffer (92.5 mM NaCl, 2.5 mM KCl, 1 mM MgCl2•6H2O, 1 mM CaCl2•2H2O, 5 mM HEPES, pH adjusted to 7.5 using NaOH) containing 0.1% BSA and 1.5 µM atropine sulfate. BSA was used to prevent peptides from sticking to plastics in the experimental apparatus, and atropine sulfate was used to block responses from potential endogenously expressed muscarinic receptors ^50–53^. Owing to reduced OR2 solubility for some peptides, all peptides were initially solubilized in 100% DMSO for 5 min, then diluted with OR2 to the desired concentration, resulting in a final DMSO concentration of 2% in all peptide solutions. For each nAChR subtype used in this manuscript, 2% DMSO did not affect peak responses evoked by the ACh EC90 (data not shown). Solutions were made fresh daily.

It is well appreciated that varying cRNA subunit injection ratios can result in expression of a mixture of nAChR isoforms. Based on the data in ^19,46^, where each subtype/isoform is expressed separately, the effective concentration 90 (EC90) was calculated (α4β2α5 12 µM; (α4β2)2β2 18 µM; concatenated β3α6β2α4β2 26 µM; α6/α3β2β3 40 µM; (α3β2)2β2 79 µM; (α3β4)2β4 220 µM; (α3β4)2α3 410 µM; (α4β2)2α4 1.16 mM; α7 1.30 mM; and (α3β2)2α3 3.05 mM).

The potential subtype selectivity of each tested peptide was determined using an nAChR subtype screening TEVC protocol. Table 1 displays the peptide sequence information. For each tested peptide, the EC90 ACh was applied for 1s to individual oocytes expressing the intended nAChR. Following an 84s OR2 wash, 100 µM peptide was pre-applied for 30s, followed by a 1s ACh EC90 application. The peptide-altered response was normalized to the initial ACh EC90 response without peptide pre-application to determine the percent inhibition caused by the peptide. Due to a priming effect of α7 nAChRs, two ACh EC90 hits were performed prior to inhibition with the peptides, and peptide-inhibited responses were normalized to the second ACh EC90 response.

For peptides that showed high α7 nAChR subtype selectivity, the apparent potency was determined. Using α7 nAChR expressing oocytes, ACh EC90 was applied twice, with 225s OR2 wash in between each application, as this subtype requires pre-exposure to ACh before the full response is evoked. Following, increasing concentrations of each peptide (0.01 - 300 µM) were pre-applied for 30s, followed by 1s of 1300 µM (EC90) ACh, followed by 225s of OR2 recording buffer in between each drug application. Responses were normalized to the second ACh EC90, peptide naïve evoked response, and the resulting data were fit using an unconstrained monophasic sigmoidal inhibition equation ([Inhibitor] vs. response - variable slope (four parameters)) with GraphPad Prism v.10 (Boston, MA) software.

To characterize the specific class of positive allosteric modulation that ARA mediates, comparative experiments using the known nAChR positive allosteric modulator (PAM) PNU- 120596 were performed. PNU-120596 has been proven to act as a type II PAM on α7 nAChRs by recovering desensitized receptors. To confirm this, experiments were performed where the α7 ACh EC90 (1300 µM) was co-applied with PNU-120596 following either 30s of PNU-120596 only application, or 30s of ACh EC90 application to reach steady state receptor desensitization. The same protocol was followed, replacing 3 µM PNU with 0.3 µM ARA to determine if ARA exhibited the same type of PAM activity as PNU-120596.

We observed that higher concentrations of ARA (≥ 10 µM) produced an outward current when applied to α7 nAChR-expressing oocytes (but not when using uninjected oocytes). The presence of this outward current precluded us from analyzing the response data using the net-charge response as used by others ^54,55^. Experiments were performed with the known nAChR antagonist MLA in an effort to block these currents. An MLA concentration of 10 nM was determined to be required to overcome ACh stimulation. Experiments were then performed where either 10 nM MLA or 100 µM ARA was pre-applied for 10s, followed by 10s of co-application. These experiments were also performed with PNU-120596 (3 µM) or Meca (10 µM) in place of MLA. We then assessed if the outward current was due to activation of a signaling cascade that involved the α7 nAChR metabotropic activities ^56–58^. Initially, we attempted to express α7(345 - 348A) nAChRs, a mutant which does not interact with Gαq and Gβγ ^57^, but the receptors did not express or were not functional in oocytes. Finally, we attempted to rectify the outward current by utilizing the compound YM-254890, which is known to inhibit the G-protein activity of the α7 nAChR ^58^. As YM-254890 needs time to penetrate cells, we added 2 - 20 µM to the oocyte buffer solution for 1 - 24hr before applying 100 µM ARA for 15 seconds.

To characterize the ARA peptide’s mechanism of antagonism, we generated ARA and ACh co-application concentration-response profiles. This was accomplished by co-applying increasing concentrations of ACh (0.010 µM - 10 mM) with a single peptide concentration (0 µM, 10 µM or 100 µM) for 1s with an 84s wash of OR2 recording buffer in between each drug application. Following the co-application responses, two applications of Imax ACh (10 mM) without peptide were performed. The first and second ACh-only responses were comparable in amplitude, indicating the washout time was sufficient to remove any residual ARA peptide. Co-application data was normalized to the second ACh Imax response. The resulting data were fit using a monophasic sigmoidal equation ([Agonist] vs. response - Variable slope (four parameters)) using GraphPad Prism v. 10 software.

TEVC experiments were conducted on at least two batches of cRNA synthesis and at least three oocyte isolations from individual frogs. The number of experimental replicates from individuals is indicated by N and followed by the number of individual oocytes (n).

### Expression of α7 nAChRs in cultured N2a cells

Mouse neuroblastoma N2a cells were purchased from the American Tissue Culture Collection (ATCC, Manassas, VA). N2a cells were maintained in EMEM supplemented with 10% FBS and 1X Penicillin-Streptomycin at 37°C and 5% CO2 in a humidified cell culture incubator. Cells were sub-cultured onto either poly-D-lysine-coated glass coverslips (MatTek, MA) or 48- well plates (VWR, Radnor, PA) and transiently transfected when dishes were approximately 70% confluent with cells. The Lipofectamine 2000 transfection reagent was used for transient transfection, according to the manufacturer’s protocol. The mammalian α7 nAChR-coding plasmid DNA and the NACHO chaperone plasmid DNA were combined at a 4:1 ratio in Opti-MEM for α7 nAChR-positive groups. We have previously successfully used and proven this technique to transfect N2a cells with α7 nAChRs, in O’Brien et al. (2024) ^19^. As N2a cells endogenously express low levels of α7 nAChR, we knocked down expression by transfecting 100 pmol of α7 siRNA (ThermoFisher) using Lipofectamine 2000 (as described above) and used these cells as our α7 nAChR knockdown control.

### N2a corrected total cell fluorescence analysis

To determine corrected total cell fluorescence (CTCF) with ImageJ (National Institutes of Health, Bethesda, MD), we calculated the mean integrated density of fluorescence in each group, using non-deconvoluted static images. To reduce bias, the observer was blind to the fluorescence channels and free-hand traced cells that appeared healthy in size and shape in the phase channel before measuring fluorescence. Area, mean, and integrated density were measured in the fluorescent channel. Three distinct circular ROIs surrounding each selected cell were drawn to measure background fluorescence. Using these values, CTCF was calculated according to the following formula: CTCF = integrated density – (area of cell x mean background fluorescence) ^59^. This process was repeated for four cell passages and transfections (N) with 30 - 50 cells (n) for each N.

### Cell viability assay

To determine if the novel peptides were cytotoxic, we assayed cell viability using the alamarBlue Cell Viability reagent, as we have previously performed ^46^. Briefly, N2a cells were seeded on 48-well plates, and α7 nAChR-positive groups were transfected as described in the *Expression of α7 nAChRs in cultured N2a cells* section. Twenty-four hours after transfection, cells were exposed to different concentrations of RVG, α-btx lp2, or ARA (0.03 - 100 µM) and incubated for an additional 24hr. 15% DMSO was added to control wells in lieu of peptide to serve as a positive control for cell death. The media was changed after 24hr, and the alamarBlue Cell Viability reagent was added to all wells at 1/10^th^ of the volume, according to the manufacturer’s instructions. After 3hr, a Tecan Spark Multimode Microplate Reader (Tecan U.S., Inc., Morrisville, NC) was used to measure changes in fluorescence (excitation: 555 nm, emission: 600 nm, bandwidth: 20 nm). Fluorescence was measured by taking 8x8 measurements per well, while keeping a 1,500 μm border to the well walls, to reduce possible detection of fluorescence transmitted from neighboring wells or interference of well walls. For cell viability experiments, large N denotes the number of 48-well plates, whereas small n designates the number of wells.

### Fluorescent confocal microscopy

To verify the positive transfection of α7 nAChR DNA, 80 nM Alexa Fluor 647-conjugated α-btx (α-btx-AF647, ThermoFisher) was added to EMEM growth medium with FBS and antibiotics 24hr post-transfection and incubated for an additional 24hr before being imaged. To determine if the RVG or ARA peptides targeted α7 nAChRs expressed in the plasma membrane of N2a cells, 30 µM FITC-tagged RVG or FITC-tagged ARA was added to the growth medium 24hr post-α7 nAChR DNA transfection and incubated for an additional 24hr. Peptide concentrations used were based on their functional potency as presented in *The α-bungarotoxin/RVG chimeric peptide, ARA, is highly selective for the α7 nAChR subtype,* results section. Cells were then rinsed three times with PBS before being imaged on the Olympus (Center Valley, PA) Fluoview FV10i Laser Scanning Confocal Microscope (AF647: 653 nm excitation, 668 nm emission wavelengths; FITC: 495 nm excitation, 519 nm emission wavelengths). Captured images were used for visualization and fluorescence CTCF quantification as described in the *N2a corrected total cell fluorescence analysis* section.

### Confocal Image Processing

After capture, images used for visualization only were deconvoluted using ImageJ and the DeconvolutionLab 2 and PSF Generator plug-ins (Biomedical Imaging Group, École polytechnique fédérale de Lausanne). Point spread functions (PSFs) were calculated using the PSF Generator and the microscope settings as described in the above sections. Using the calculated PSFs and the DeconvolutionLab2 plug-in, images were deconvoluted using the Richardson-Lucy algorithm with 10 iterations ^60^.

To determine if α-btx-AF647 or the ARA peptide is internalized into cells, we generated three-dimensional representations of cells labeled with either construct. Single or small clusters of visually healthy, labeled cells were chosen for 3D visualization and imaged at the above-mentioned microscope settings with a 1 µm slice size. The image stacks of each channel were first separated from one another, then deconvoluted separately and developed into 3D projections using the 3D project function with interpolation. Phase and fluorescent channels were then combined into one image to create a multi-channel 3D model. Internalization of RVG-FITC has been previously published in O’Brien et al. (2024) ^19^.

### Data Analysis

All quantified data were analyzed using GraphPad Prism v.10. Data sets comparing two groups were analyzed using Welch’s *t*-tests. For data sets with three or more groups, a One or Two-Way ANOVA was performed based on the number of independent variables. Data are displayed as the mean ± S.D. Throughout all statistical analyses, significance levels are denoted as follows: ^ns^P > 0.05, *P < 0.05, **P < 0.01, ***P < 0.001, and ****P < 0.0001.

## Results

### The α-bungarotoxin/RVG chimeric peptide, ARA, is highly selective for the α7 nAChR subtype

To begin designing an RVG-derived CPP that is α7 nAChR target-selective, we explored combining regions of α-btx lp2 with RVG. We first separated the RVG peptide and α-btx lp2 peptide into distinct regions based on protein sequences (Table 1). The RVG peptide contains a sequence in the middle with no homology to the α-btx lp2 peptide, so the RVG peptide was broken into three regions (R1, R2, R3) while the α-btx lp2 peptide was broken into only two regions (R1, R3). Using TEVC electrophysiology, we determined the apparent potency for each set of synthetic α-btx lp2 and RVG fragments by pre-applying increasing concentrations of each peptide (0.01 - 300 µM) for 30s, followed by a 1s ACh EC90 (1300 µM) stimulation. The full-length α-btx lp2 peptide was the most potent (2.5 µM (Confidence Interval (CI) 2.0, 3.1 µM)), followed by α-btx R1 (59 µM (CI 50, 69 µM)) and α-btx R3 (462 µM (CI 333, 981 µM)) (Figure 1A). The RVG peptide had an enhanced apparent potency (37 µM (CI 28, 42 µM)) when compared to RVG R1, RVG R2, or RVG R3 peptides. RVG R3 was the most potent (176 µM (CI 147, 213 µM)) of the three fragments, with RVG R1 and RVG R2 having minimal or no actions on α7 nAChR function (Figure 1B).

**Figure 1.**
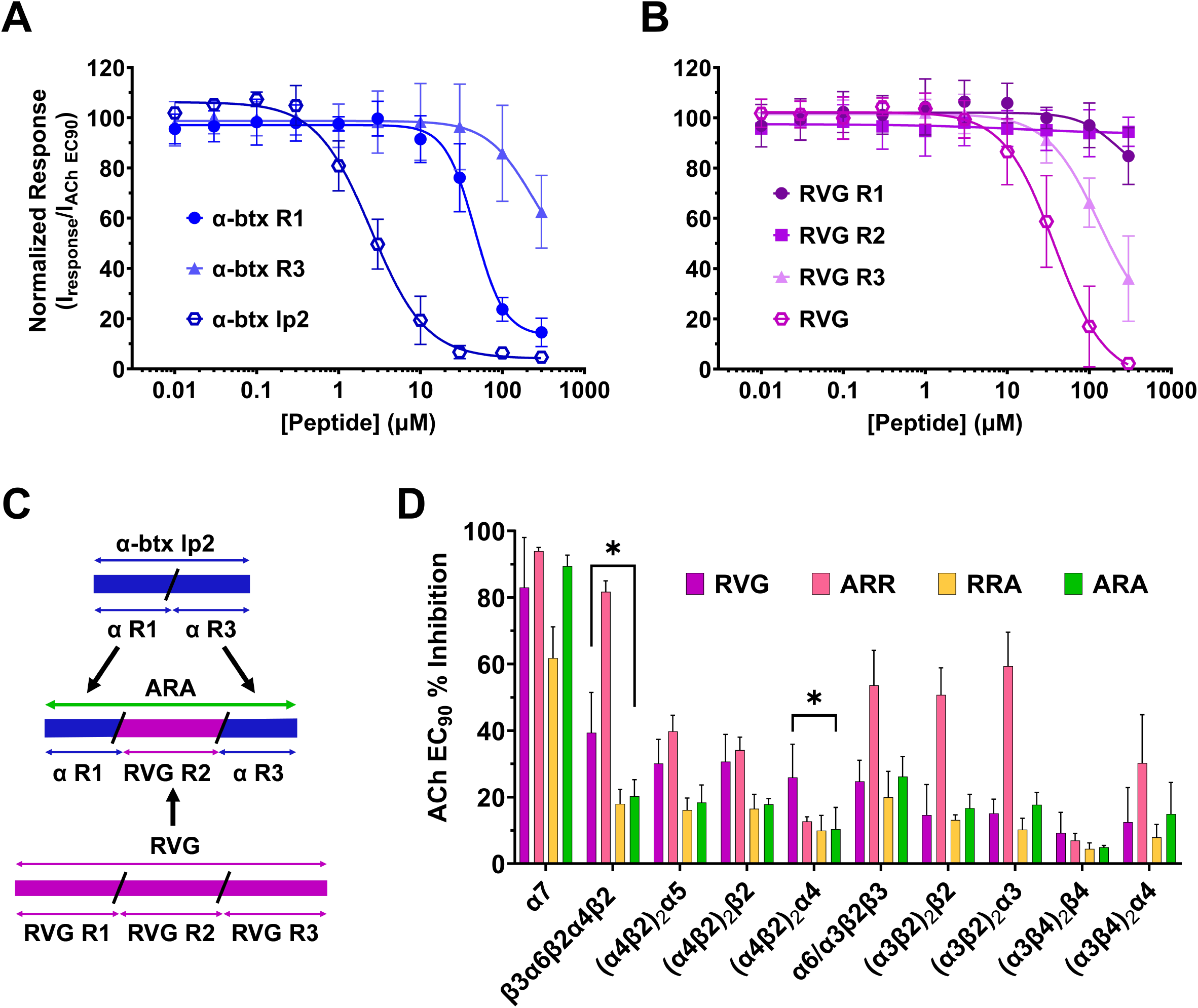
Chimeric peptides generated from regions of RVG and α-btx lp2 have differing selectivity profiles. TEVC analysis of concentration-dependent effects of the test peptides on nAChR-expressing oocytes. **(A)** α-btx lp2 is highly potent (IC50 = 2.5 µM (CI 2.0, 3.1 µM) on the α7 nAChR (N = 3 - 4, n = 5 - 7). **(B)** RVG components R1, R2, and R3 are not very potent on the α7 subtype (N = 3 - 4, n = 5 - 8). **(C)** Schematic illustration of chimeric peptide design and nomenclature. **(D)** In comparison to the RVG, ARA increases inhibition of the α7 nAChR (83 ± 15% to 89 ± 3%) while significantly decreasing inhibition on the β3α6β2α4β2 (*P = 0.0145) and the (α4β2)2α4 receptor (*P = 0.0482, Two-Way ANOVA with Tukey’s multiple comparison test) (N = 3 - 5, n = 3 - 9). Points are the mean ± S.D.

We then combined the regions of RVG and α-btx lp2 to construct an α7 nAChR-selective peptide with the goal of maintaining the CPP properties of RVG while enhancing target selectivity (Figure 1C). We hypothesized that the peptide, which contained α-btx R1, RVG R2, and RVG R3 (ARR), would likely be the most effective at antagonizing α7 nAChRs, as the α-btx R1 and RVG R3 had the most independent inhibitory actions.

The first step in evaluating whether we could generate a potent α7 nAChR subtype selective chimeric peptide was to understand the selectivity of the RVG parent peptide. To this end, the peptide was screened on nAChR subtypes present in the CNS, including α7, α4β2, α3β2, α3β4, concatenated β3α6β2α4β2, α6/α3β2β3, and α4α5β2 ^61,62^. Each isoform of the α4β2, α3β2, and α3β4 subtypes was tested separately for a total of 10 neuronal nAChRs. The RVG peptide is known to antagonize the nAChR response to ACh by competing for the orthosteric binding pocket^19^. We performed a subtype selectivity screen, which allowed us to quickly determine if the peptides were efficacious on the evaluated nAChRs (Figure 1D). Using TEVC, clamped oocytes expressing the desired nAChR were first exposed to their specific ACh EC90 (α4β2α5 12 µM; (α4β2)2β2 18 µM; β3α6β2α4β2 26 µM; α6/α3β2β3 40 µM; (α3β2)2β2 79 µM; (α3β4)2β4 220 µM; (α3β4)2α3 410 µM; (α4β2)2α4 1.16 mM; α7 1.29 mM; (α3β2)2α3 3.05 mM) to provide a baseline response. Following washout of ACh, oocytes were pre-exposed to 30s of 100 µM peptide followed by an ACh EC90 stimulation. This protocol allowed us to quickly determine how much the RVG peptide inhibited each nAChR subtype.

The RVG parent peptide inhibits the ACh response of α7 nAChR the most (83 ± 15%) of any of the tested nAChRs (Figure 1D). The α4β2 subtype is the most abundant nAChR expressed in the CNS ^63,64^ and both isoforms of α4β2 (HS 27 ± 10%, LS 22 ± 7%), as well as other α4β2- containing subtypes (α4α5β2 30 ± 7% and β3α6β2α4β2 39 ± 12%) ACh-evoked responses were also inhibited by RVG. The remaining subtype ACh mediated currents were inhibited < 26% (α6/α3β2β3 25 ± 6%; (α3β2)2β2 15 ± 9%; (α3β2)2α3 15 ± 4%; (α3β4)2β4 7 ± 3%; (α3β4)2α3 12 ± 10%). These results indicate that the RVG peptide inhibits the α7 subtype the most, a key feature we attempted to enhance with our designed chimeric peptides, but also antagonizes the function of other abundantly CNS-expressed nAChRs.

To reduce the probability of off-target effects, we aimed to generate a CPP with high α7 nAChR selectivity by incorporating components of α-btx. To screen the three RVG/α-btx chimeric peptides, we used the same protocol as described above for the RVG peptide. Peptide ARR, which is composed of α-btx R1, RVG R2, and RVG R3, maximally inhibited the α7 nAChR ACh-mediated currents (94 ± 1%) (Figure 1D). However, it also displayed the least selectivity, as five of the 10 nAChRs tested had > 50% inhibition (Figure 1D). The RRA (RVG R1, RVG R2, and α- btx R3) peptide inhibited heteromeric nAChRs ACh-evoked currents < 20% (Figure 1D), but only inhibited the α7 nAChR ACh currents by 62 ± 9% (Figure 1D). The remaining ARA (α-btx R1, RVG R2, and α-btx R3) peptide robustly inhibited α7 nAChR ACh currents (89 ± 3%), while minimally inhibiting the heteromeric subtypes ≤ 18% (α6/α3β2β3 18 ± 5%; (α3β2)2β2 17 ± 4%; (α3β2)2α3 18 ± 4%; (α3β4)2β4 4.9 ± 0.6%; (α3β4)2α3 15 ± 10%). Importantly, the ARA peptide reduced antagonization of the α4β2 containing subtypes (HS 18 ± 2%, LS 10 ± 7%, α4α5β2 20 ± 5%, β3α6β2α4β2 26 ± 6%), demonstrating a significant improvement for α7 nAChR selectivity compared to the RVG peptide (β3α6β2α4β2 *P = 0.0145 and (α4β2)2α4 *P = 0.0482, Two-Way ANOVA with Tukey’s multiple comparison test).

### The ARA peptide has an improved apparent potency and efficacy

To determine if the apparent potency of the ARA peptide was enhanced compared to the RVG peptide, as suggested by the selectivity screen, we generated concentration-response profiles using TEVC electrophysiology (Figure 2). As described for RVG and α-btx lp2 (Figures 1A and B), concentration-response profiles were generated using α7 nAChR expressing oocytes and 30s of pre-applied increasing concentrations of the ARA peptide (0.01 - 300 µM) before ACh EC90 (1300 µM) stimulation. Responses were compared to the peptide naïve ACh-evoked response to normalize the data. The ARA peptide had a significantly enhanced apparent potency (19 µM (CI 16, 22 µM)) compared to the RVG peptide (37 µM (CI 28, 42 µM)) (*P = 0.0102), but was not as potent as the α-btx lp2 peptide (2.5 µM (CI 2.0, 3.1 µM)) (***P = 0.0004, One-Way ANOVA with Dunnett’s multiple comparison test) (Figure 2, Supplemental Table 1).

**Figure 2.**
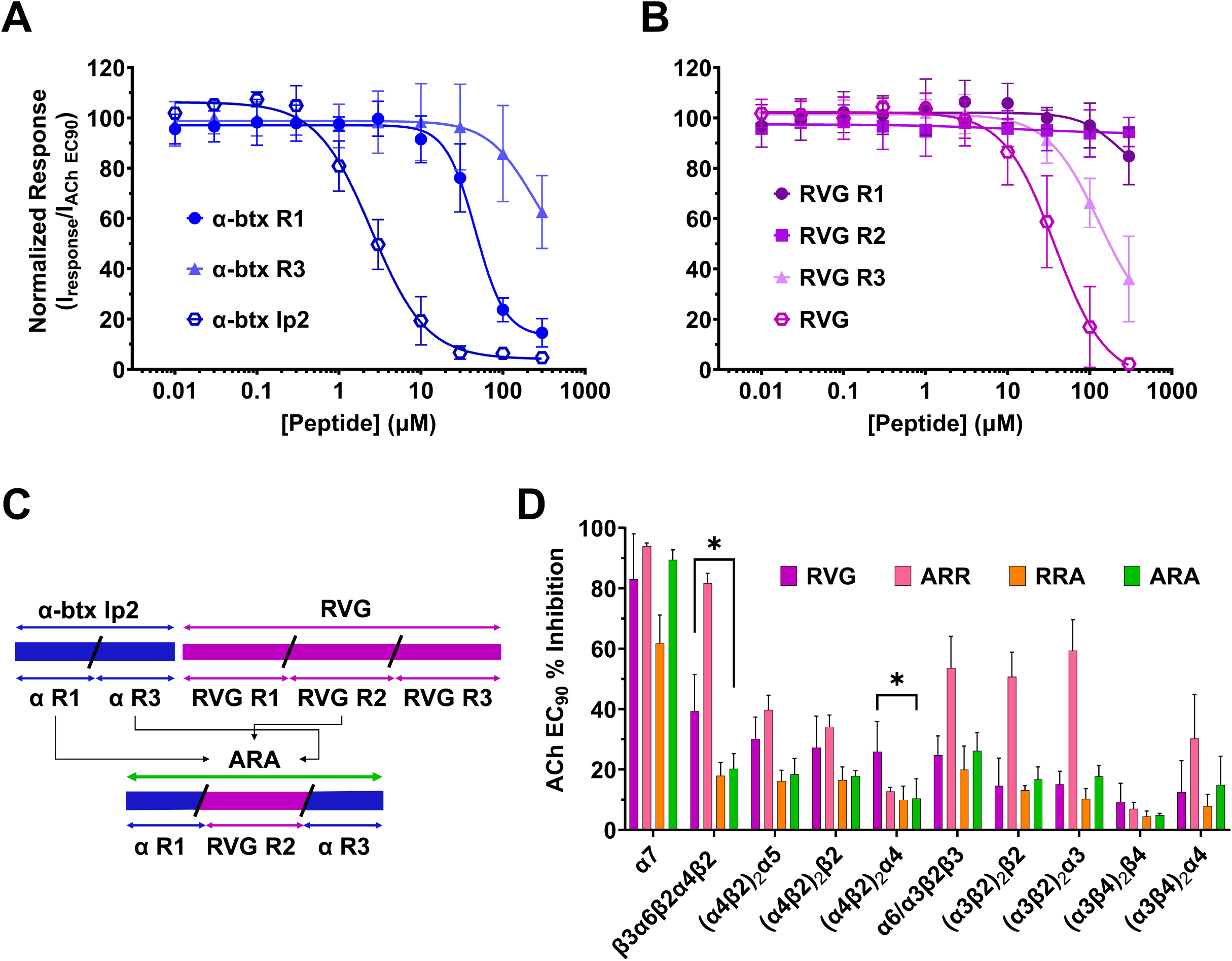
The concentration-response profile for ARA displays an increase in the apparent potency for the α7 nAChR compared to the RVG peptide. Concentration-response profiles of the peptides’ antagonistic effects on α7 nAChR ACh-evoked responses were generated using TEVC electrophysiology. The ARA peptide had a significantly enhanced apparent potency (19 µM (CI 16, 22 µM)) compared to the RVG peptide (37 µM (CI 28, 42 µM)) (*P = 0.0102), but was not as potent as the α-btx lp2 peptide (2.5 µM (CI 2.0, 3.1 µM)) (***P = 0.0004, One-Way ANOVA with Dunnett’s multiple comparison test). Points are the mean ± S.D. (N = 3 - 4, n = 5 - 8).

In Figure 2, the low concentrations of ARA slightly enhanced (114 ± 7%) the ACh EC90 response, suggesting that ARA may function as a PAM. Type I PAMs increase the agonist-induced peak current, and type II PAMs, in addition to enhancing the peak response, also slow the rate of receptor desensitization ^65^. To investigate if ARA is a type I or type II PAM, we compared a peptide naïve ACh response to one where 0.3 µM was pre-applied for 30s, which resulted in an increased peak response (130 ± 10%, Figure 3A). In 2011, Collins et al. conducted a series of experiments demonstrating that PNU-120596 is an α7 nAChR type II PAM that prolongs the receptors’ return to baseline and can reactivate desensitized receptors ^66^. Unlike PNU-120596, with application of ARA, there was no significant change in response shape or time to return to baseline. When 0.3 µM ARA was applied to α7 nAChRs during ACh (1300 µM) induced steady-state, no recovery of desensitized receptors was observed (Figure 3B). This data indicates there may be some slight type I PAM activity of ARA at low concentrations.

**Figure 3.**
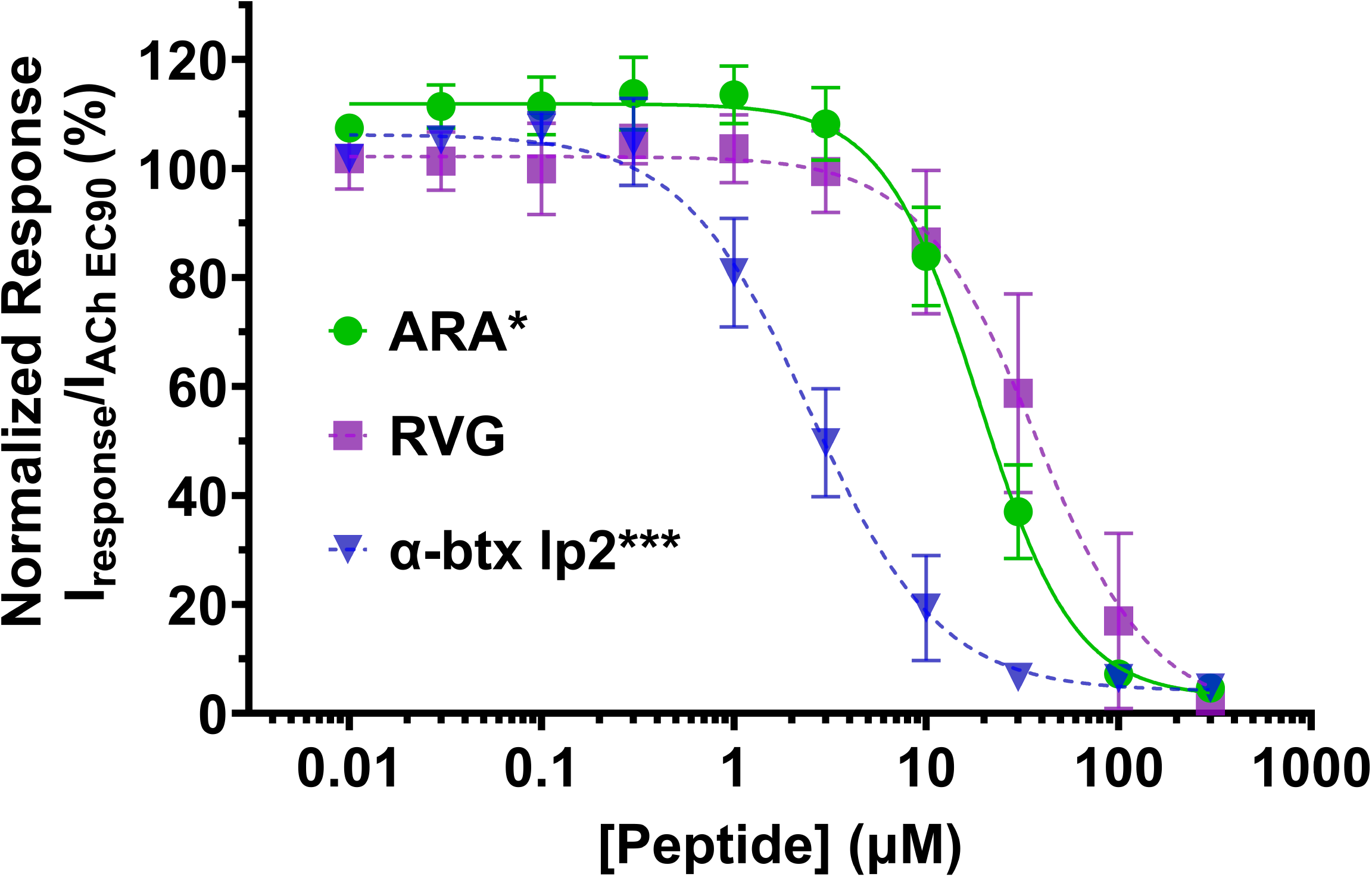
Low concentrations of ARA potentiate ACh responses, but ARA cannot re- activate desensitized α7 nAChRs. α7 nAChR-expressing oocytes were either exposed to 0.3 µM ARA prior to or after ACh application, and responses were recorded using TEVC electrophysiology. **(A)** Representative trace of the α7 subtype illustrating potentiation of an ACh-induced response following preapplication of 0.3 µM ARA. Experiments were repeated N = 4, n = 5. **(B)** ARA is not capable of reactivating desensitized α7 receptors (example trace shown, repeated N = 3, n = 6).

Interestingly, at high ARA concentrations (10 - 300 µM), we observed a small outward current (3 - 15 nA) when the peptide was applied to α7 nAChR expression oocytes (Figure 4A). This current was absent in oocytes not expressing α7 nAChRs (Figure 4B), demonstrating that it was α7 nAChR-mediated. This outward current, coupled with the slight PAM activity (Figure 3A), suggested to us that the ARA peptide may have silent agonist properties.

**Figure 4.**
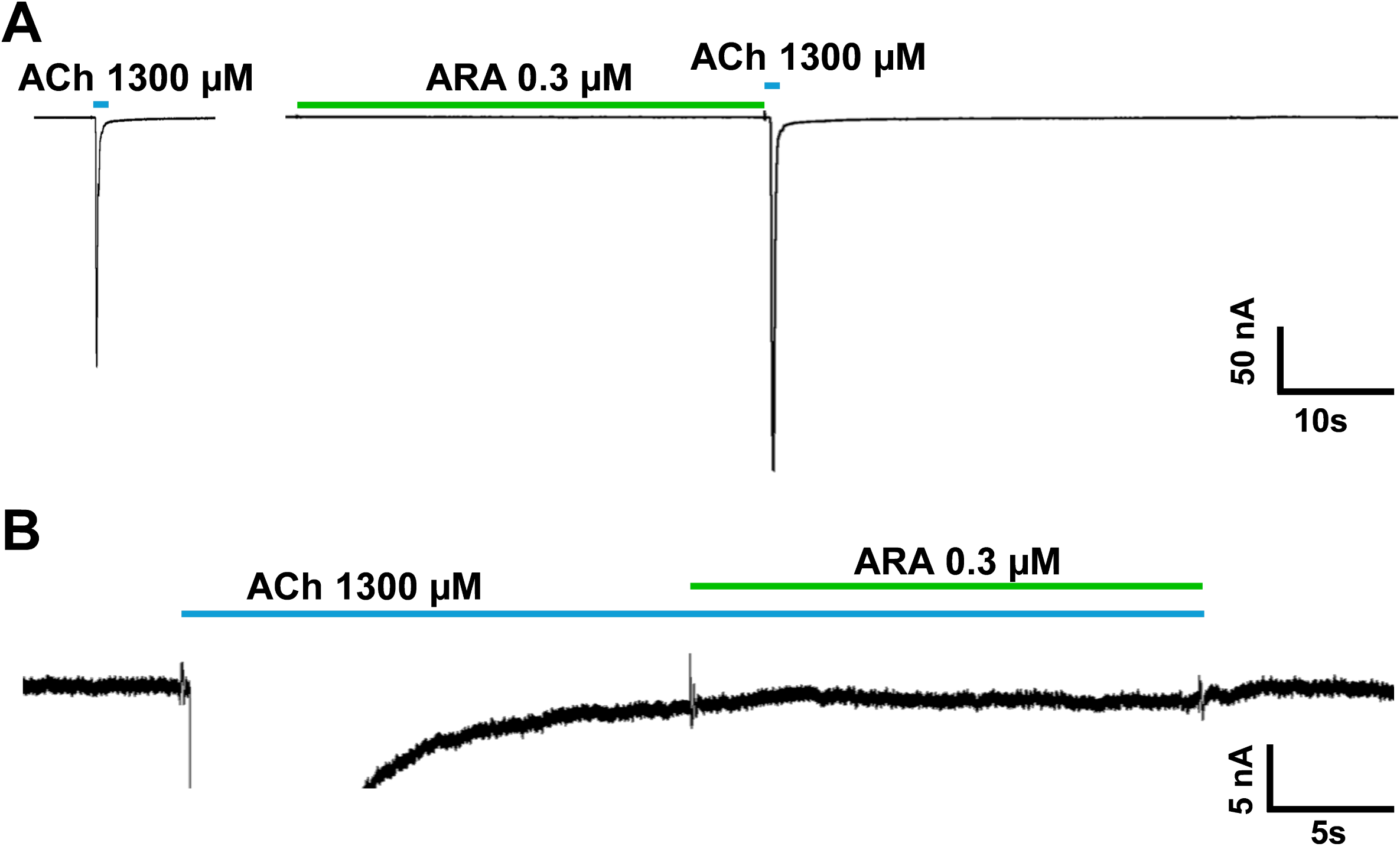
High concentrations of ARA evoke an outward current that is dependent on the α7 nAChR. α7 nAChR or uninjected oocyte representative responses to ARA, PNU-120596, MLA, and Meca were captured using TEVC electrophysiology. **(A)** High concentrations of ARA produce an outward current in α7 nAChR-expressing oocytes **(B)** that is not present without α7 nAChR expression (example traces shown, repeated N = 3, n = 3). The outward current generated by high ARA concentration is unaffected by **(C)** the α7 nAChR PAM PNU-120596 (example trace shown, repeated N = 3, n = 7), **(D)** α7 nAChR competitive antagonist MLA (example trace shown, repeated N = 3, n = 5), or the **(E)** non-competitive channel blocker Meca (example trace shown, repeated N = 3, n = 6).

Silent agonists are a class of compounds that, upon binding, induce a conformational change that stabilizes the desensitized state rather than an open state. Previous work with silent agonist NS6740 has shown that using a type II PAM, such as PNU-120596, can shift the α7 nAChR towards a conducting state without activation ^67,68^. To test if ARA has silent agonist activity at higher concentrations, we pre-applied either ARA (100 µM) or PNU-120596 (3 µM) for 10s, followed by a co-application of ARA with PNU-120596 for another 10s (Figure 4C). No receptor activation was seen in either instance, and the outward current created by ARA was unaffected. We next tried to block the outward current with α7 nAChR competitive antagonist MLA or the non-competitive antagonist Meca. Neither pre- nor co-application with either antagonist could mitigate the small outward current created by application of 100 µM ARA (Figure 4D and E).

In addition to ionotropic activity, the α7 subtype has been shown to activate metabotropic responses by directly interacting with intracellular G-proteins ^69,70^. Given that the ARA-evoked current is α7-mediated (Figure 4A and B) and the current generated is slow relative to an ACh response, we reasoned that ARA may trigger activation of the G-protein to activate a distinct effluxing ion channel. We attempted to express the α7(345 - 348A) mutant, which lacks the G-protein binding capacity ^57^, in oocytes. Unfortunately, with injecting 20 - 120 ng of cRNA and waiting 3 - 14 days post-injection, we were unable to observe responses to ACh (Supplemental Figure 1A). Next, we attempted to inhibit the endogenous G-protein in oocytes using YM-254890, a known plasma membrane permeable inhibitor of the Gα protein in the Gq/11-protein family ^71,72^. We bath-applied 2 - 20 µM YM-254890 for 1 - 24hr, and regardless of concentration or incubation time, we were unable to prevent or reduce the outward current generated by ARA (Supplemental Figure 1B). Future work will include structural ARA modification to eliminate the outward current.

### ARA is a competitive antagonist

An nAChR competitive antagonist competes with ACh for the orthosteric binding pocket and functionally reduces the ACh apparent potency without changing efficacy ^45,73–75^. As we have previously determined that RVG is a competitive antagonist, we hypothesized that ARA antagonizes α7 nAChRs by the same mechanism ^19^. To this effort, we co-applied increasing concentrations of ACh (1 µM - 10 mM) with 0 µM, 10 µM, or 100 µM ARA and normalized each response to a 10 mM ACh response without ARA present (Figure 5). Applying either 10 or 100 µM ARA resulted in a rightward shift of the concentration response curve compared to ACh without ARA co-application (+ 0 µM ARA IC50: 208 µM (CI 181, 240 µM), + 10 µM ARA: 322 µM (CI 271, 390 µM), and + 100 µM ARA: 518 µM (CI 445, 611 µM), Supplemental Table 2). While significant changes in the apparent potency were observed (+ 10 µM ARA *P = 0.0204 and + 100 µM ARA ****P < 0.0001, One-Way ANOVA with Dunnett’s multiple comparison test), there was no change in efficacy with ARA co-application. These data are consistent with ARA inhibiting α7 nAChRs via competitive antagonism.

**Figure 5.**
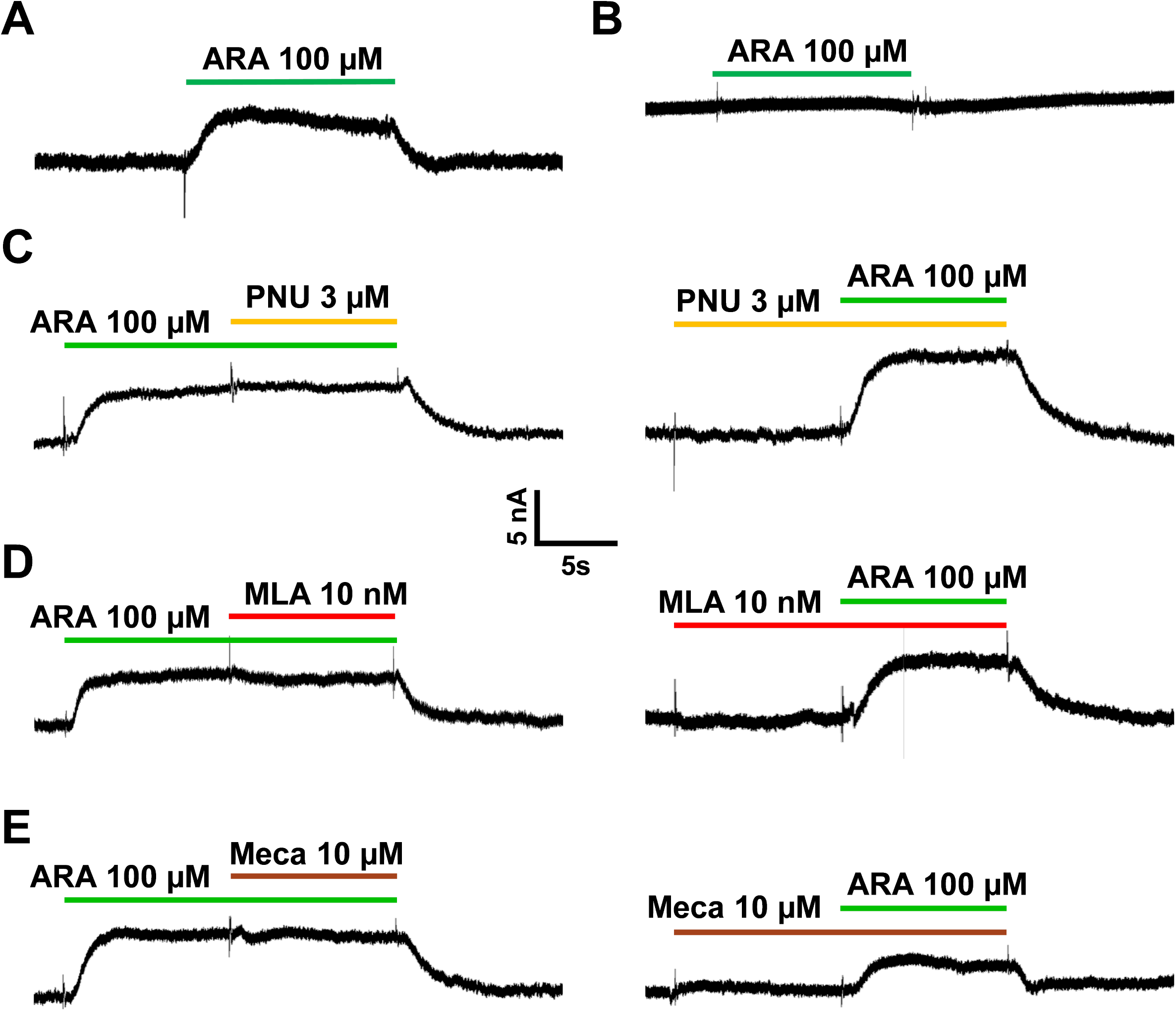
ARA competes with ACh for binding to the α7 nAChR. α7 nAChR-expressing oocytes were co-exposed to increasing concentrations of ACh without or with ARA, and responses were observed using TEVC electrophysiology. Co-application of ARA with increasing concentrations of ACh significantly shifts the response curve to the right, resulting in significant decreases in the apparent potency (10 µM ARA *P = 0.0246, 100 µM ARA ****P < 0.0001, One-way ANOVA with Dunnett’s multiple comparison test) without affecting efficacy. Points are the mean ± S.D. (N = 3 - 4, n = 8 - 13).

### ARA is not cytotoxic

Peptide therapies are attractive, partially due to their low cytotoxicity profile ^76^. However, a subset of CPPs, specifically those designed to selectively target and eliminate cancerous cells, are cytotoxic ^77,78^. In efforts to develop ARA as a potential drug-delivery CPP, we performed a comprehensive cytotoxicity screening using N2a cells transfected with the α7 subunit DNA as a preliminary look at the potential safety profile. We used alamarBlue cell viability reagent and found that after 24hr of incubation with 0.03 - 100 µM of RVG, α-btx lp2, or ARA peptides, no cytotoxic effects were observed (^ns^P > 0.05, One-way ANOVA with Tukey’s multiple comparison test) (Figure 6 and Supplemental Figure 2). A 15% DMSO control was used to demonstrate positive detection of the dead cells (****P < 0.0001, One-way ANOVA with Dunnett’s multiple comparison test, untreated group as control).

**Figure 6:**
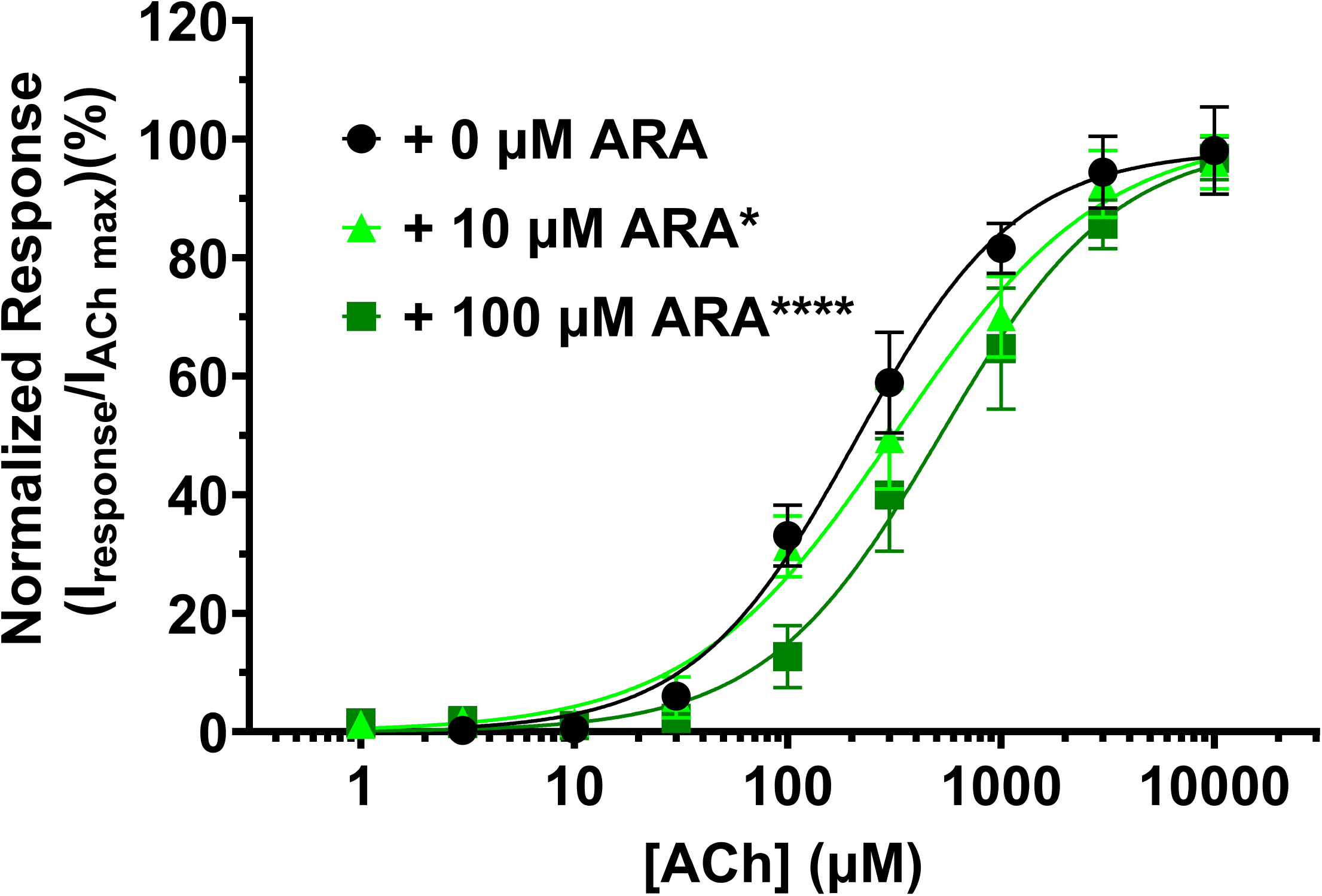
RVG, α-btx lp2, and ARA peptides are not cytotoxic at functional concentrations. Cytotoxicity profiles for **(A)** the RVG parent peptide, **(B)** the α-btx lp2 peptide, and **(C)** ARA using α7 nAChR-transfected N2a cells. Peptides (0.03 - 100 μM) that were pre-applied for 24hr and assessed by the alamarBlue Cell Viability Assay show no significant cytotoxic effects (One-way ANOVA with Tukey’s multiple comparison test, ****P < 0.0001). Points are the mean ± S.D. (N = 3, n = 6 - 9).

### ARA preferentially labels N2a cells expressing α7 nAChRs

Some drug delivery studies have shown that RVG derivatives, such as RVG29, are capable of delivering cargo, including molecules and siRNAs, into the CNS and only require concentrations in the high nanomolar to low micromolar range ^12,14^. We next aimed to determine if ARA selectively targets only α7 nAChRs by using the mammalian N2a cell line with a neuronal phenotype. Each peptide was tagged with a fluorophore: ARA was labeled with FITC (ARA-FITC), full-length α-btx was conjugated with AF648 (α-btx-AF648), and RVG was bound to FITC (RVG- FITC). α7 nAChR-over expressing N2a cells were incubated for 24hr with either α-btx-AF647, RVG-FITC, or ARA-FITC peptides.

Live-cell confocal microscopy showed robust α-btx-AF648 (80 nM) binding to α7 nAChR DNA-transfected cells (Figure 7A and D). Conversely, non-transfected cells displayed minimal labeling (Figure 7B) relative to non-labeled cells (Figure 7D). To verify that the slight increase in labeling using non-transfected cells could be attributed to endogenous α7 nAChR expression, cells were transfected with α7 siRNA and treated with α-btx-AF647 24hr later. α7 siRNA-treated cells had less α-btx-AF647 labeling (Figure 7C) than non-transfected treated cells, and were not different from untreated cells (Figure 7D). Quantification of fluorescence as CTCF (Figure 7E) verifies the highest α-btx-AF647 binding in α7 nAChR DNA-transfected cells (386.7 ± 39.8 %, ****P < 0.0001, One-way ANOVA with Tukey’s multiple comparison test), while non-transfected and siRNA-transfected cells show only low levels of CTCF (151.6 ± 43.9 %, ^ns^P = 0.1298, and 117.6 ± 7.2 %, ^ns^P = 0.8, respectively), comparable to background fluorescence levels in untreated cells (100 ± 11.7 %). This emphasizes that the binding of α-btx-AF647 was due to the presence of α7 nAChRs.

**Figure 7:**
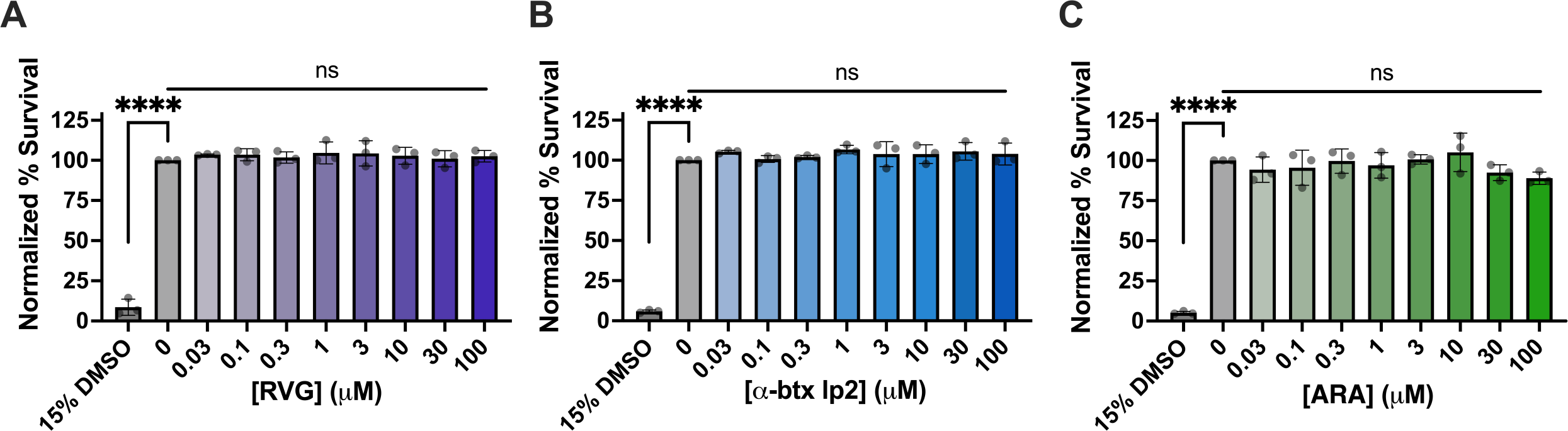
Demonstration of α7 nAChR-specific labeling with α-btx-AF647. Confocal images of **(A)** α7 nAChR-transfected, **(B)** non-transfected, and **(C)** α7 nAChR siRNA-transfected N2a cells treated with 80 nM α-btx-AF746 for 24hr show selective binding of α-btx-AF647 to cells overexpressing α7 nAChRs and only little binding to cells with residual endogenous expression. **(D)** Untreated N2a cells were imaged in the AF647 fluorescent channel. Α-btx-AF647 selectively interacts with cells expressing α7 nAChRs and interacts little with cells not transfected with α7 nAChRs. **(A’ - D’)** Same as (A-D) without the phase channel. Quantification of CTCF **(E)** demonstrates high fluorescence in cells expressing α7 nAChRs, with no significant fluorescence in non-transfected and α7 nAChR siRNA-transfected cells (One-way ANOVA with Tukey’s multiple comparison test, N = 4, n = 120 - 174). Values are the mean ± S.D. and were normalized to untreated controls **(D)**.

As shown in Figure 1D, RVG appears to be mildly selective for α7 nAChR. However, the rabies virus glycoprotein targets other host proteins, including the neural cell adhesion molecule ^20,21^, p75 neurotrophin receptor ^23^, metabotropic glutamate receptor subtype 2 ^25^, and integrin β1 ^26^, which are all expressed in N2a cells. We thus anticipated that the application of RVG-FITC would result in the labeling of the above-mentioned RVG targets, in addition to α7 nAChRs. α7 nAChR over-expressing N2a cells were treated with 30 µM of RVG-FITC, and the resulting fluorescence was visualized using live-cell confocal microscopy. Cells over-expressing α7 nAChRs showed the highest RVG-FITC-associated fluorescence (Figure 8A). However, cells that were not α7 nAChR transfected (Figure 8B) or had been treated with α7 siRNA (Figure 8C) showed elevated levels of fluorescence as well, compared to non-treated cells, demonstrating substantial non-α7 nAChR binding of the RVG-FITC peptide (Figure 8D). Quantification of RVG- FITC CTCF (Figure 8E) confirmed a significant increase in fluorescence in α7 nAChR-transfected cells relative to blank cells, nearly tripling the CTCF value (290.7 ± 34.5 %, ****P < 0.0001, One-way ANOVA with Tukey’s multiple comparison test). In contrast, non-transfected (213.4 ± 34.9 %, **P = 0.0056) and α7 siRNA-transfected (191.1 ± 56.7 %, *P = 0.0237) cells exhibited less pronounced increases in CTCF, but still significantly more than background (100 ± 11.4 %). These results align with previous findings, substantiating that RVG utilizes nAChRs for interaction with cells ^19,27,28^, but does not solely rely on the α7 nAChR subtype.

**Figure 8:**
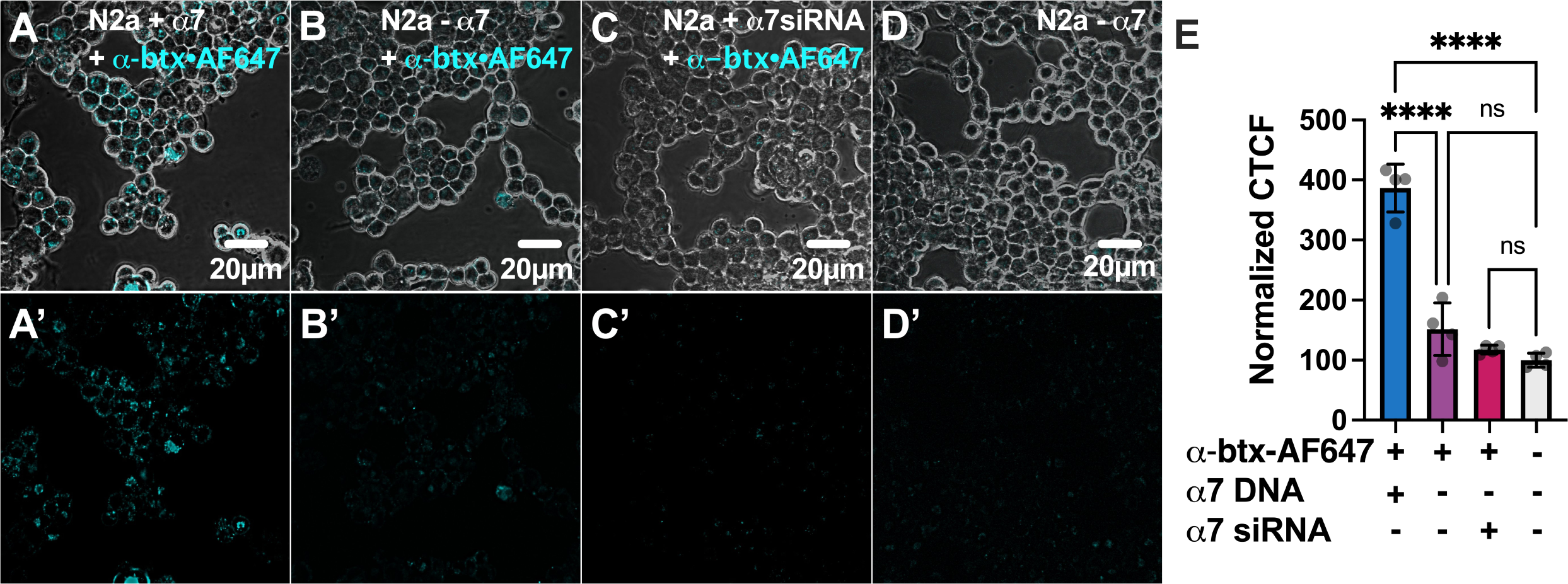
The RVG CPP non-selectively interacts with N2a cells. Confocal images of **(A)** α7 nAChR-transfected, **(B)** non-transfected, and **(C)** α7 nAChR siRNA-transfected N2a cells treated with 30 μM RVG-FITC for 24hr showing RVG binding to N2a cells regardless of α7 nAChR presence. **(D)** Untreated N2a cells imaged in the FITC fluorescent channel. RVG interaction with N2a cells is not dependent on the expression of the α7 nAChR. **(A’ - D’)** Same as (A-D) without the phase channel. Quantification of CTCF **(E)** confirms similar fluorescence levels in cells with and without α7 nAChRs (One-way ANOVA with Tukey’s multiple comparison test, N = 4, n = 120 - 135). Values are the mean ± S.D. and were normalized to untreated controls **(D)**.

As the ARA peptide is a chimeric peptide with components of the α7 nAChR-selective α- btx and the non-selective RVG peptide, we tested ARA selectivity using α7 nAChR over- expressing N2a cells as performed for the two parent peptides. The ARA-FITC fluorescence labeling was visualized by live-cell confocal microscopy. Cells expressing the α7 nAChR exhibited pronounced labeling with 30 µM ARA-FITC 24hr post-exposure (Figure 9A). In contrast, non-transfected cells displayed minimal fluorescence (Figure 9B), and siRNA knockdown of endogenous α7 nAChRs diminished the ARA-FITC labeling (Figure 9C) as seen in non-transfected, non-treated groups (Figure 9D). These data support that α7 nAChR was responsible for the robust ARA-FITC labeling. Quantitative analysis of fluorescence intensity (Figure 9E) revealed a substantial increase in ARA-FITC-associated fluorescence in α7 nAChR-transfected cells (312.3 ± 93.2 %, **P = 0.0011, One-way ANOVA with Tukey’s multiple comparison test) in comparison to blank cells (100 ± 11.4 %). ARA-FITC labeling was drastically reduced in non-transfected cells (137.7 ± 66.0 %, ^ns^P = 0.8) and α7 siRNA-treated cells (127.6 ± 16.8 %, ^ns^P = 0.9), which were not significantly different from untreated cells.

**Figure 9:**
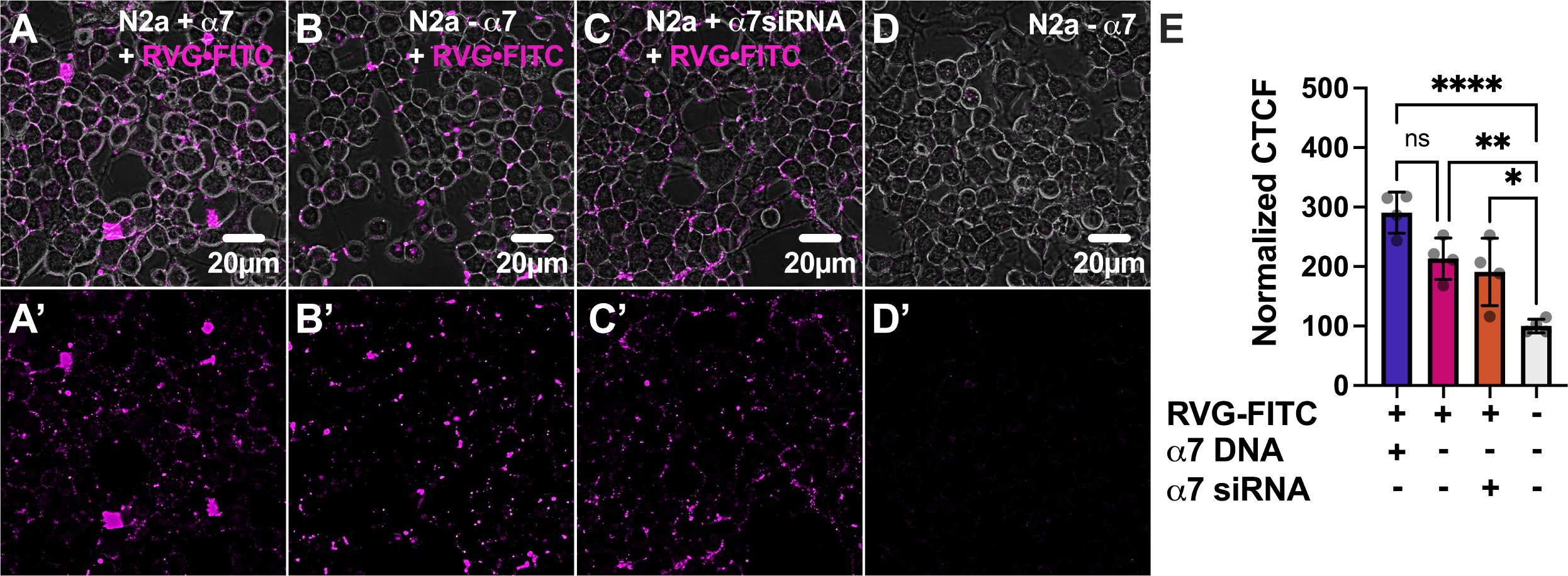
ARA preferentially interacts with cells expressing α7 nAChRs. Confocal images of. **(A)** α7 nAChR-transfected, **(B)** non-transfected, and **(C)** α7 nAChR siRNA-transfected N2a cells treated with 30 μM ARA-FITC for 24hr demonstrate high fluorescence levels associated with ARA- FITC in cells expressing α7 nAChRs. **(D)** Untreated N2a cells were imaged in the FITC fluorescent channel to account for autofluorescence. Endogenously expressed α7 nAChRs in N2a cells contribute to the staining in non-transfected N2a cells, which is reduced by transfection of α7 siRNA. **(A ’- D’)** Same as (A-D) without the phase channel. Quantification of CTCF **(E)** demonstrates significantly more fluorescence in cells expressing α7 nAChRs (One-way ANOVA with Tukey’s multiple comparison test, N = 4, n = 117 - 126). Values are the mean ± S.D. and were normalized to the untreated control **(D)**.

### The ARA peptide is a CPP

Prior studies have demonstrated the ability of RVG-derived peptides to penetrate cultured cells, neurons, and even the brains of mice and rats by a receptor-mediated mechanism ^12,14,19^. Our chimeric peptide, ARA, incorporates segments of the neurotoxin α-btx, introducing a novel structural aspect that could hinder cellular entry.

To assist with visualization of the location of the fluorescence, N2a cells expressing α7 nAChRs were treated with α-btx-AF647 (40 nM) or ARA-FITC (30 µM) for 24hr prior to imaging. Z-stack images were captured and compiled to create 3D projects (Figure 10 and Supplemental Videos 1 and 2). Upon careful inspection of the live-cell confocal images, we observed very different fluorescence staining patterns of α-btx-AF647 and ARA-FITC (Figure 10). α-btx-AF647 fluorescence occurred mostly around the perimeter of the cells (Figure 10A). In comparison, ARA- FITC fluorescence was located throughout the interior, demonstrating that our ARA peptide was inside the cells (Figure 10B). It has been shown for multiple other RVG derivatives that nAChRs are crucial interaction scaffolds for these peptides to connect to the cellular plasma membranes and use these receptors as vehicles to internalize, likely through receptor-mediated endocytosis ^13,79,80^. Given the retention of some RVG structures within ARA, it appears that the ARA peptide employs α7 nAChRs to enter the cell by a receptor-mediated process.

**Figure 10:**
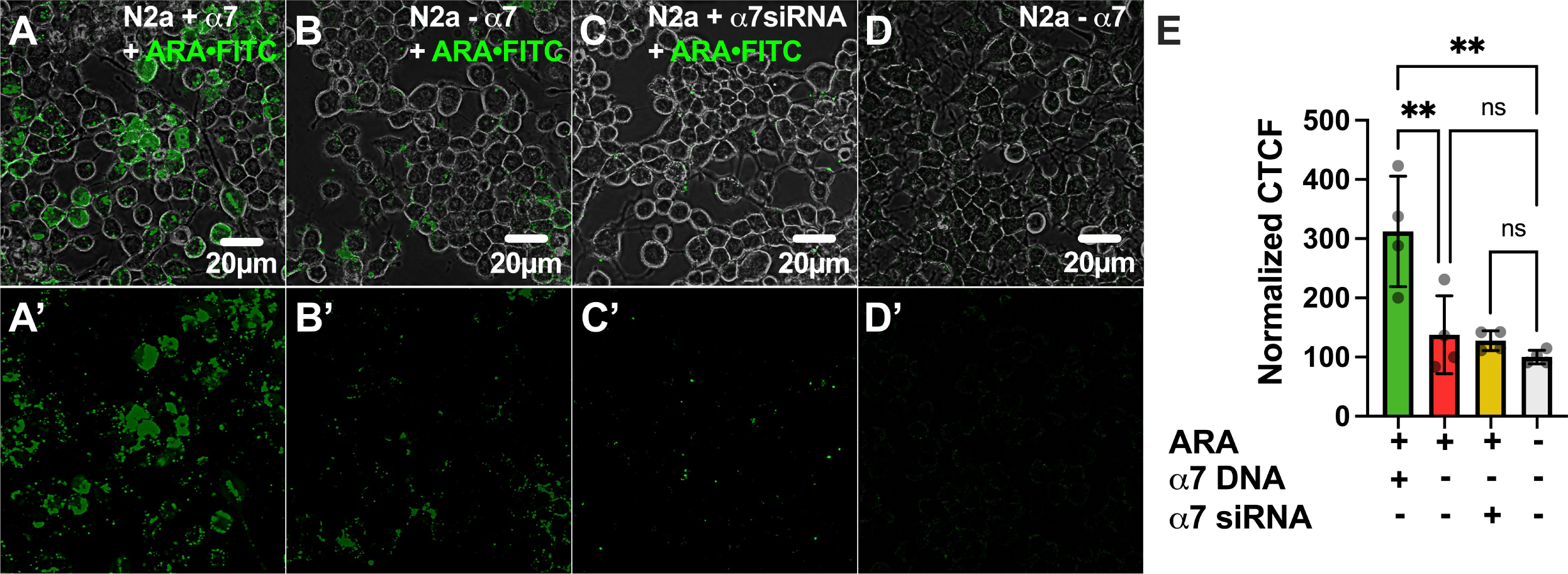
ARA transports FITC cargo into α7 nAChR-expressing cells. Representative 3D projections from z-stack confocal images of α7 nAChR-transfected N2a cells (z-stack slice size: 1 μm), treated with **(A)** 30 μM FITC-tagged ARA peptide or **(B)** 40 nM α-btx-AF647 for 24hr prior to imaging. **(A’ and B’)** Rotated around the y-axis by 40°. ARA appears in the cell interior, while α-btx binds to the cell surface without internalization.Table

## Discussion

In this study, we developed and characterized a novel chimeric CPP created through combining the regions of α-btx responsible for α7 nAChR targeting and the rabies virus glycoprotein region hypothesized to mediate cell entry. ARA is highly selective for the α7 nAChR subtype, unlike the parent RVG peptide, with a good apparent potency in the low micromolar concentration (Figures 1 and 2). ARA has multiple functional modalities on the α7 subtype. At low concentrations (< 3 µM), ARA potentiates ACh-induced currents a modest amount. At higher concentrations (> 3 µM), ARA is a competitive antagonist, attenuating the ACh-induced currents and shifting the ACh apparent potency to the right (Figure 5). Curiously, > 10 µM of ARA not only reduced the ACh response but also induced a slow outward current that cannot be removed with other antagonists (Figure 4). Attempts to determine if the outward current generated by ARA application was due to activation of a G-protein-mediated signaling cascade were unsuccessful. Future work to improve ARA will include structural refinements to remove the production of an outward current.

Using α7 nAChR-expressing N2a cells, a mouse-derived cell line with a neuronal phenotype, we performed an initial toxicity screen and found that ARA is not cytotoxic up to the highest concentration tested (100 µM) (Figure 6). We demonstrate a clear α7 nAChR dependence for the cellular interaction of ARA in both our electrophysiology (Figure 1) and cell culture assays (Figure 9). Using cultured cells, ARA-FITC substantially labeled cells expressing α7 nAChRs, with nominal labeling observed in cells with endogenous α7 nAChR expression (Figure 9). This suggests that the ARA peptide requires the presence of α7 nAChRs on the cell surface to bind to N2a cells. To further accentuate the importance of α7 nAChRs for cellular interaction of ARA, we used α7 nAChR-siRNA to knock down receptor expression. Cells subjected to α7 nAChR knockdown exhibited the least amount of fluorescent labeling in response to peptide treatment (Figure 9E). Comparison to the α7 nAChR-selective antagonist α-btx revealed analogous interaction profiles, highlighting the enhanced selectivity of ARA over the less-selective RVG peptide (Figures 7 and 8). These findings further establish that our ARA chimeric peptide exhibits enhanced selectivity for α7 nAChRs as targets on neuronal-like cells, displaying minimal non-selective binding *in vitro* in comparison to the RVG parent peptide. Visual observations were substantiated by quantitative analysis of fluorescence values associated with FITC tags, providing compelling evidence that α7 nAChRs are crucial for the interaction of ARA with neuronal-like cells. RVG29, a CPP that can transport therapeutic cargo into the mouse brain when given peripherally, has proven not to be cytotoxic, and multiple doses given over weeks are well tolerated ^12,14,81^. Given that ARA is more target-selective than RVG (Figure 1D), it is plausible that concentrations required for *in vivo* drug delivery may be even lower than what is currently being used with RVG29. The lack of cytotoxic effects positions ARA as a promising candidate for further investigation in drug delivery, highlighting its potential for safer and more effective therapeutic interventions.

One of our most exciting findings is that ARA is an α7 nAChR target-selective CPP. Visualization through 3D projections revealed fluorescence in the intracellular space, indicating that, like RVG29, ARA shares the capability to enter and transport cargo, including FITC, into cells (Figure 10B). Receptor-mediated endocytosis, as proposed for RVG29, is a potential mechanism for how ARA gains access to the inside of the cell ^13,79,80^. It remains to be determined the cargo size limitation ARA can transport across the cell membrane, and if the cargo can escape the endosome.

Non-selective CPPs have been used to transport cargo across *in vitro* models of the BBB and into the brain *in vivo* using mice ^82–84^. RVG29, attached to nine unnatural arginine amino acids (9dR), has been used to transport several types of siRNA across the mouse BBB when injected into the tail vein ^12,14^. However, RVG peptides have been demonstrated not to be target-directed, which limits their therapeutic potential by generating off-target effects (Figures 1 and 8 ^19^). A commonly overlooked feature of drug delivery systems is the functional effect of the CPP on the cellular target(s). ARA has multiple modalities on α7 nAChRs, including potentiation and antagonism, depending on the concentration. Future work developing ARA as an α7 nAChR-selective therapeutic delivery system could have applications for the treatment of neurological conditions involving the α7 nAChR, including Alzheimer’s disease and major depressive disorder 39,85–87.

The development of CPPs as drug-delivery tools is a significant advancement in enabling the transport of therapeutic cargo into the brain. CPPs offer a promising strategy for overcoming the BBB, which has long been a challenge in treating neurological disorders. Here we have presented the generation of ARA, a novel, potent, and highly target-selective α7-nAChR CPP. CPPs typically have lower binding affinities than antibodies but less potential for immunogenicity, susceptibility to proteases due to their limited size (< 30 amino acid residues), stronger penetration ability, easier synthesis and modification, and lower production costs ^88,89^. These positive attributes of peptides make CPPs active therapeutic agents.

## Conclusions

Clinically relevant applications of ARA include attaching an siRNA 9dR hook, or cargo-loaded nanoparticles or exosomes, as has been accomplished using RVG29 ^12,14,90,91^. These modifications allow for the potential improvement in delivering a variety of therapeutics to cells expressing α7 nAChRs. Critically, high target selectivity of ARA ensures that the therapeutic payload is delivered to specific cell types or tissues, enhancing the therapeutic efficacy while minimizing the off-target effects. For drug delivery systems to be effective and safe, they must exhibit low cytotoxicity to avoid damaging healthy cells and tissues, a property that ARA may also possess based on our cell assays. The combination of these factors makes ARA an exciting and promising approach for advancing targeted therapies. The presented data serve as a foundation for future investigations, including *in vivo* testing, to validate and further explore the potential of ARA as an efficient drug delivery agent for intracellular and CNS-targeted applications.

## Acknowledgments

The authors would like to thank James Janoso for his technical microscopy expertise and the Center for Transformative Research in Metabolism (National Institute of General Medical Sciences of the National Institutes of Health under Award Number P20GM130443) for supporting the Microscopy Core at the University of Alaska Fairbanks.

## Ethics approval

Ecocyte Bioscience, an IACUC-certified company, provided the oocytes used in the electrophysiology studies. The mouse N2a cell line was purchased from ATCC.

## Declaration of interest

The authors declare no competing interests.

## Author contributions

All authors have contributed significantly and agree with the manuscript’s content. MMW is responsible for the original conception and design. BOB, LW, and MMW performed the data analysis and data interpretation. All authors participated in drafting the paper and revising it critically for intellectual content. Final approval of the manuscript was performed by MMW. All authors meet the criteria for authorship as per the ICMJE guidelines.

## Data availability

The data presented in this study is available on request from the corresponding author.

## Additional information

### Funding

Research reported in this publication was supported by the National Institute of Mental Health of the National Institutes of Health (MMW by award number R03MH135358). The authors received funding from an Institutional Development Award (IDeA) from the National Institute of General Medical Sciences of the National Institutes of Health (grant number P20GM103395) (MMW, BOB, and LW), Alaska Center for Innovation, Commercialization, and Entrepreneurship (grant from the Office of Naval Research, #N00014-18-1-2725) (MMW), and the Higher Education Investment Fund, provided by the state of Alaska through the University of Alaska (LW and BOB).

---

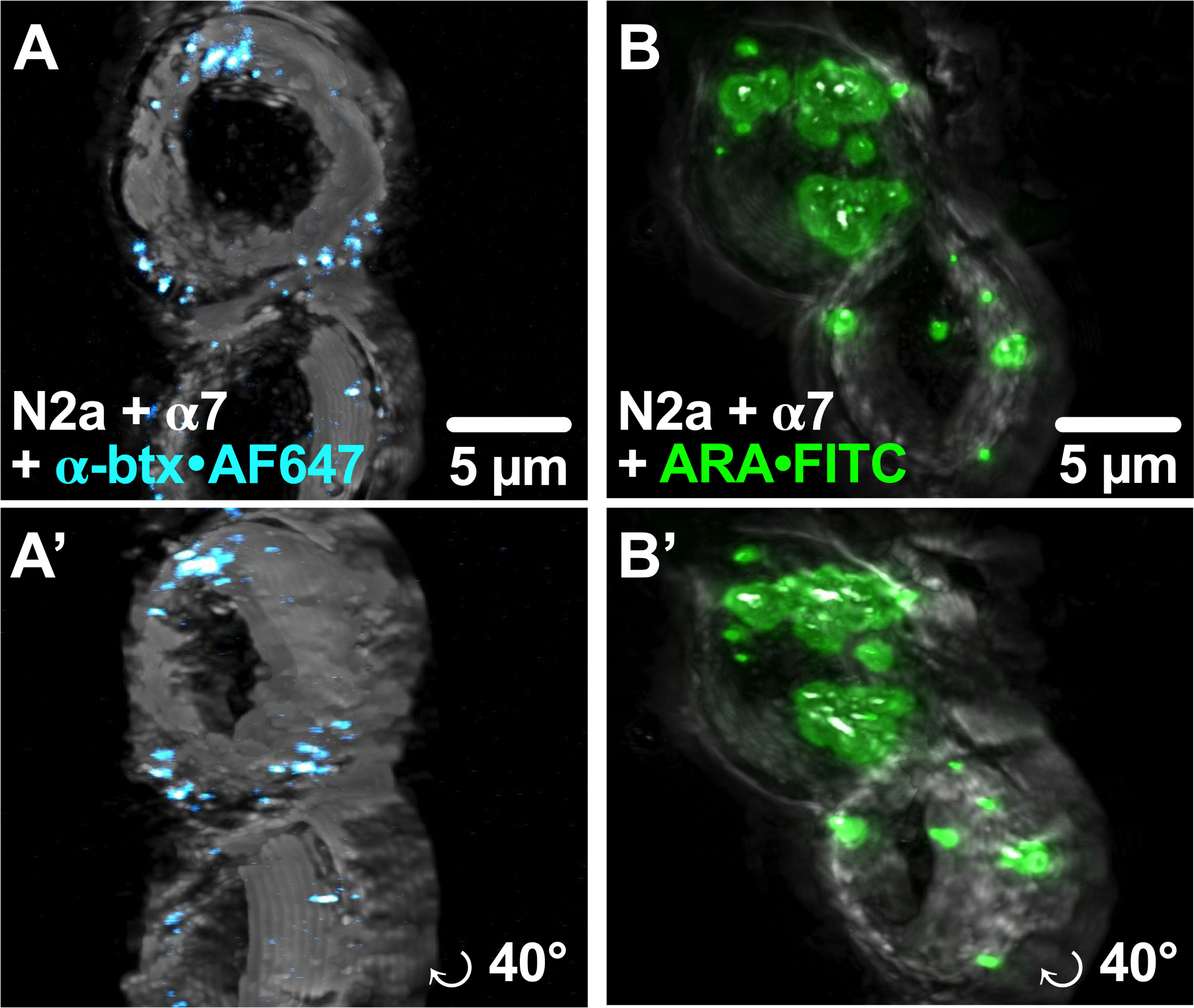

## Notes

### Competing Interest Statement

The authors have declared no competing interest.

