## Supplemental Information for "Development of a novel alpha7-nicotinic acetylcholine receptor-selective cell-penetrating peptide for intracellular cargo transport"

**Supplemental Table 1. Quantified data for the peptide pre-application α7 nAChR inhibition concentration-response profiles.** Pre-application peptide concentration response data were fit using non-linear curve fitting. RVG R1 and R2 potency and $N_H$ values could not be determined (N.D.) as these peptides minimally inhibited the α7 subtype. Data are presented in Figures 1 and 2.

| Peptide Treatment | N (n) | $IC_{50}$ (μM) (95% CI) | $N_H$ (95% CI) |
|---|---|---|---|
| α-btx R1 | 4 (6) | 59 (50 – 69) | -1.6 (-1.3 to -2.1) |
| α-btx R3 | 4 (7) | 462 (333 – 981) | -1.3 (-0.7 to -2.6) |
| α-btx lp 2 | 3 (5) | 2.5 (2.0 – 3.1) | -1.2 (-1.0 to -1.4) |
| RVG R1 | 4 (7) | N.D. | N.D. |
| RVG R2 | 4 (6) | N.D. | N.D. |
| RVG R3 | 3 (5) | 176 (147 – 213) | -1.2 (-0.9 to -1.4) |
| RVG | 4 (8) | 37 (28 – 42) | -1.5 (-1.2 to -1.9) |
| ARA | 4 (5) | 19 (16 – 22) | -1.7 (-1.4 to -2.0) |

**Supplemental Table 2. Calculated parameters for ARA competitive antagonist Ach concentration-response profiles.** Non-linear curve fitting was used to analyze the data, and are shown in Figure 5.

| Co-application ACh concentration response profiles | | | |
|---|---|---|---|
| **Peptide Treatment** | **N (n)** | **$EC_{50}$ (μM) (95% CI)** | **$N_H$ (95% CI)** |
| none | 3 (8) | 208 (181 – 240) | 1.14 (1.01 – 1.29) |
| 10 μM ARA | 4 (12) | 322 (271 – 390) | 0.90 (0.81 – 1.00) |
| 100 μM ARA | 4 (13) | 518 (445 – 611) | 1.06 (0.95 – 1.18) |

**Supplement Figure 1: The ARA outward current could not be prevented by blocking metabotropic G-protein signaling. (A)** Example trace of an oocyte injected with 40 ng of α7(345 - 348A) nAChR cRNA and tested five days post-injection exhibited no response to 1300 μM ACh application. Similar experiments testing a range of cRNA concentrations and recording from 3 - 14 days post-cRNA injection all resulted in no ACh-evoked responses (N = 4, n = 12). **(B)** Example trace of an α7 nAChR expressing oocyte soaked in 20 μM YM-254890 for 2hr, and exposed to 100 μM ARA still produced an outward current (N = 2, n = 5).

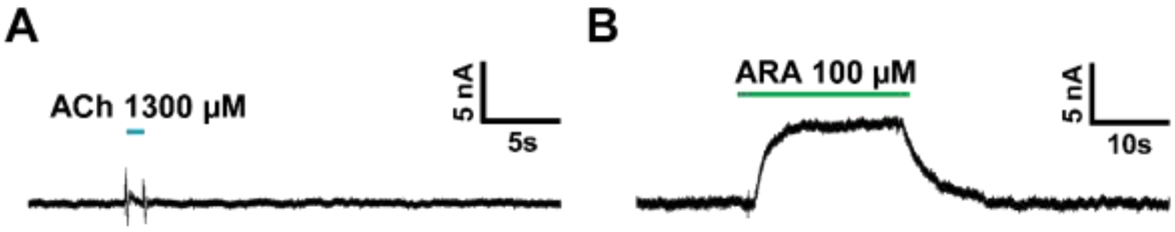

**Supplement Figure 2: RVG, α-btx lp2, and ARA show no cytotoxicity at functional concentrations in non-transfected cells.** Cytotoxicity profiles for **(A)** the RVG parent peptide, **(B)** the α-btx lp2 peptide, and **(C)** ARA on non-transfected N2a cells assessed by the alamarBlue Cell Viability Assay (One-way ANOVA with Tukey's multiple comparison test, ****P < 0.0001). Points are the mean ± S.D. (N = 3, n = 6 - 9).

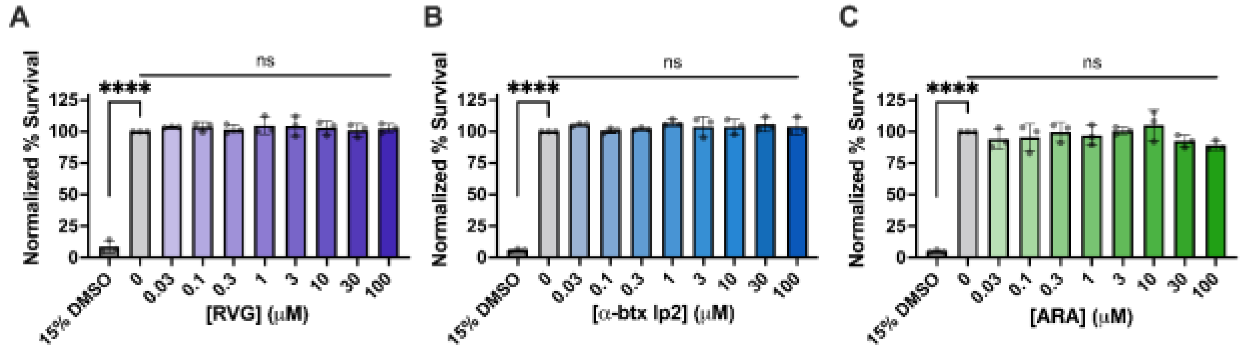

**Supplement Video 1:** 3D movie animation of α-btx-AF647-labeled N2a cells transfected with α7 nAChR and NACHO DNA, showing binding of α-btx-AF647 to the plasma membrane of cells. Note that α-btx-AF647 fluorescence is not found in the interior of the cells.

**Supplement Video 2:** 3D movie animation of ARA-FITC-labeled α7 nAChR expressing N2a cells. Strong ARA-FITC fluorescence can be inside the cells.
